## Supplementary file S1 for "Intragenomic sequence variations in the second internal transcribed spacer (ITS2) ribosomal DNA of the malaria vector *Anopheles stephensi*"

### Supplementary information

#### S1: Annotation features of GenBank entries

```
>Feature gb|MW676288
<1    52          5.8S ribosomal RNA
53    522         internal transcribed spacer 2
523   >1679       28S ribosomal RNA

>Feature gb|MW676289
<1    52          5.8S ribosomal RNA
53    520         internal transcribed spacer 2
521   >1677       28S ribosomal RNA

>Feature gb|MW676290
<1    52          5.8S ribosomal RNA
53    520         internal transcribed spacer 2
521   >1677       28S ribosomal RNA

>Feature gb|MW676291
<1    52          5.8S ribosomal RNA
53    520         internal transcribed spacer 2
521   >1677       28S ribosomal RNA

>Feature gb|MW676292
<1    52          5.8S ribosomal RNA
53    522         internal transcribed spacer 2
523   >1679       28S ribosomal RNA

>Feature gb|MW676293
<1    52          5.8S ribosomal RNA
53    522         internal transcribed spacer 2
523   >1679       28S ribosomal RNA

>Feature gb|MW676294
<1    52          5.8S ribosomal RNA
53    522         internal transcribed spacer 2
523   >1679       28S ribosomal RNA

>Feature gb|MW676295
<1    52          5.8S ribosomal RNA
53    522         internal transcribed spacer 2
523   >1679       28S ribosomal RNA

>Feature gb|MW732930
<1    52          5.8S ribosomal RNA
53    520         internal transcribed spacer 2
521   >1677       28S ribosomal RNA

>Feature gb|MW732931
<1    52          5.8S ribosomal RNA
```

| | | |
|---|---|---|
| 53 | 520 | internal transcribed spacer 2 |
| 521 | >1677 | 28S ribosomal RNA |
